# Meiosis related cancer-testis genes correlate with tumor aneuploidy and immune infiltration in 21 cancer types

**DOI:** 10.1101/608877

**Authors:** Xuewei Wang, Yide Xu, Yuting Chang, Sihan Ju, Liu Yang, Qufei Qian, Yao Chen, Shuaizhou Chen, Na Qin, Zijian Ma, Juncheng Dai, Hongxia Ma, Guangfu Jin, Erbao Zhang, Cheng Wang, Zhibin Hu

**Affiliations:** Department of Epidemiology and Biostatistics, Center for Global Health, School of Public Health, Nanjing Medical University, Nanjing 211166, China; Department of Bioinformatics, School of Basic Medical Sciences, Nanjing Medical University, Nanjing 211116, China; State Key Laboratory of Reproductive Medicine, Nanjing Medical University, Nanjing 211116, China; Jiangsu Key Lab of Cancer Biomarkers, Prevention and Treatment, Jiangsu Collaborative Innovation Center for Cancer Personalized Medicine, Nanjing Medical University, Nanjing 211116, China

**Keywords:** CT genes, aneuploidy, cancers

## Abstract

The meiosis stage of spermatogenesis and the aneuploidy generation in tumorigenesis are highly linked. Cancer-testis genes (CT genes) were specifically expressed in testis and cancers and may serve as the molecular basis of the shared features between spermatogenesis and tumorigenesis. In the present study, we integrated multi-omics data from bulk samples and cell lines, and single cell RNA-Seq data from testis and two tumors to systematically investigate the association between CT genes and aneuploidy. After ranking genes according to their association with aneuploidy level, we found that CT genes, especially CT genes specifically expressed in meiosis, showed a consistently positive correlation with aneuploidy and homologous recombination deficiency (HRD) level. Similar results were also found in the CT non-coding genes. Then, we constructed a regulatory network based on the single cell RNA-seq data and revealed that the gain of accelerator transcriptional factors (TFs) E2F7 and E2F8 and the loss of the stabilizer TFs RFX2 and NFYA aberrantly activated CT genes in the cancers and were associated with the increase of HRD level. Finally, we found that the association between CT genes and cytotoxic infiltrating lymphocytes can be influenced by the HRD level. In sum, our results revealed the evidence of pseudomeiotic functions of CT genes in the aneuploidy generation process in the cancer cells. The results may help illuminate the origin of aneuploidy in cancers and guide future immunotherapy targeting CT antigens.

## Introduction

Recent studies found the similar characters of tumorigenesis and gametogenesis, such as sustained mitosis of spermatogonia and infinite proliferation of cancer cell; haploidy in the meiosis of spermatocyte and aneuploidy in cancer cell; migration of primordial germ cell and metastasis of cancer cell(1). Thus, it was proposed that genes involved in the gametogenesis can be re-activated and consequently drive the development of cancer independently in the absence of mutations in known oncogenes and tumor suppressor genes. Cancer-testis (CT) genes, which express only in germ cells and cancer cells, may be the basis of the above theory and participated in the process of carcinogenesis and cancer metastasis. Our previous work systematically identified CT genes in cancer patients and revealed that CT genes could be a new source of epi-driver candidates of cancers (2,3), which were also supported by the other recent studies (4). Several CT genes, such as MAGEA3, FATE1, and LIN28B have been proved to contribute to the infinite proliferation of cancer cells (3–5). Besides protein coding CT genes, our previous work found that a number of non-coding genes also presented the cancer-testis expression patterns and the function of CT-lncRNA THOR and LIN28B-AS1 in cancers were also deciphered recently (2,3,6). The results mentioned above suggested that protein-coding and non-coding CT genes could be the molecular basis of proliferation similarity between cancer cells and germ cells.

In addition to the similarity of proliferation ability, both germ cells and cancer cells showed ploidy alterations of the genome: the meiosis of spermatocytes generates haploid spermatids and cancer cells carried aneuploidy (7). Aneuploidy is an unbalanced number of chromosomes and is a predominant cancer feature occurring in nearly 90% of solid tumors. The aneuploidy level in cancers is highly correlated with the reduced cytotoxic immune cells infiltration (8). However, the origin and the role of aneuploidy in tumorigenesis is still unknown. There is emerging evidence that the inappropriate activation of meiotic chromosome regulator genes results in the chromosomal aberration and genome instability. HORMAD1, one of the earliest CT genes, are essential for the meiosis. The expression of HORMAD1 in breast cancer patients can promote nonhomologous end joining, an error-prone DNA repair pathway, and therefore driving genomic instability (9). Recent TCGA study on aneuploidy also found the enrichment for the spermatogenesis pathway when investigating the aneuploidy-correlated genes (7). However, there is no study systematically investigate the correlation between CT genes and aneuploidy.

In this study, we included transcriptomic and genomic data from 8,879 TCGA patients, 777 CCLE cell lines and single cell RNA-Seq data from testis to comprehensively illuminate a strong correlation between CT genes expressed in meiosis stage and aneuploidy events in cancers and create CTatals2 to query and visualize the correlation (http://45.62.103.54:41062/ctatlas2/). As well as protein-coding CT genes, non-coding CT genes were also associated with aneuploidy in cancers. With single-cell RNA-Seq data from cancer cells, we constructed a regulatory network to illuminate that the loss of “stabilizer” transcription factor RFX2/NFYA and the gain of “accelerator” transcription factor E2F7/E2F8 in cancers contribute to the correlation between CT genes and aneuploidy. Moreover, we presented data that the immunogenicity of classic CT antigens in cancer patients may be greatly weakened because of their correlation with aneuploidy.

## Results

To systematically investigate whether CT genes were correlated with the aneuploidy level in cancer patients, we involved transcriptomic and genomic data from 21 cancer types, including bulk samples from 8,879 TCGA patients with solid tumors (Supplementary Table 1). We involved 44,460 genes which expressed in at least 1% of all samples in the following analysis. To avoid the influence of the copy number of the genes and the purity of samples, we calculated the Spearman’s partial correlation coefficients between aneuploidy score and the expression of each gene (TPM) after adjusting the gene’s copy number and the purity. A ranked list of correlation coefficients was analyzed by gene set enrichment analysis (GSEA) with the predefined CT gene lists. We conducted the GSEA analysis on the protein-coding and non-coding genes, respectively.

### A consistently positive correlation between meiosis-related CT protein-coding genes and aneuploidy score

Firstly, we defined three types of CT protein-coding genes according to the evidence of testis-specific expression, as described in our previous work. A total of Interestingly, we observed a significant enrichment of high evidence CT genes in the GSEA analysis as well as moderate and low evidence CT genes (NES_high_ = 1.76, P=1.45 × 10^-4^; NES_moderate_ = 1.57, P=4.80 × 10^-4^; NES_low_ = 1. 61, P=3.10× 10^-3^, Figure 1A). Because spermatogenesis is a tightly regulated developmental process involved multiple stages, we included the single-cell RNA-seq data (10) of 2845 testis cells and further classified CT genes into five groups according to specific expression of germ cell subtypes: spermatogonia (Spg), spermatocyte (Spc) and spermatids (Spt) specifically expressed CT genes, testis-universal (Tu) expressed CT genes and other CT genes. In the case of GSEA analysis on the five subsets of high evidence CT genes above, we observed that Spg-and Spc-specific and testis-universal CT genes were significantly enriched (NES_Spg_ = 2.46, P=1.88 × 10^-4^; NES_Spc_ = 2.14, P=1.72× 10^-4^; NES_Tu_ = 1.79, P=1.71 × 10^-4^), while Spt-specific CT genes was not (NES_Spt_ = 0.77, P=0.98)(Figure 1A). The results in the moderate and low evidence CT genes were similar (Figure 1A). To further validate our classification, we involved an independent RNA-Seq data which include the transcriptomic profile of cells from three well-defined germ cell subtypes in testis and classify CT genes accordingly. Similarly, the enrichment of Spg- and Spc-specific and testis-universal CT genes were validated (Supplementary Figure 1).

**Figure 1.**
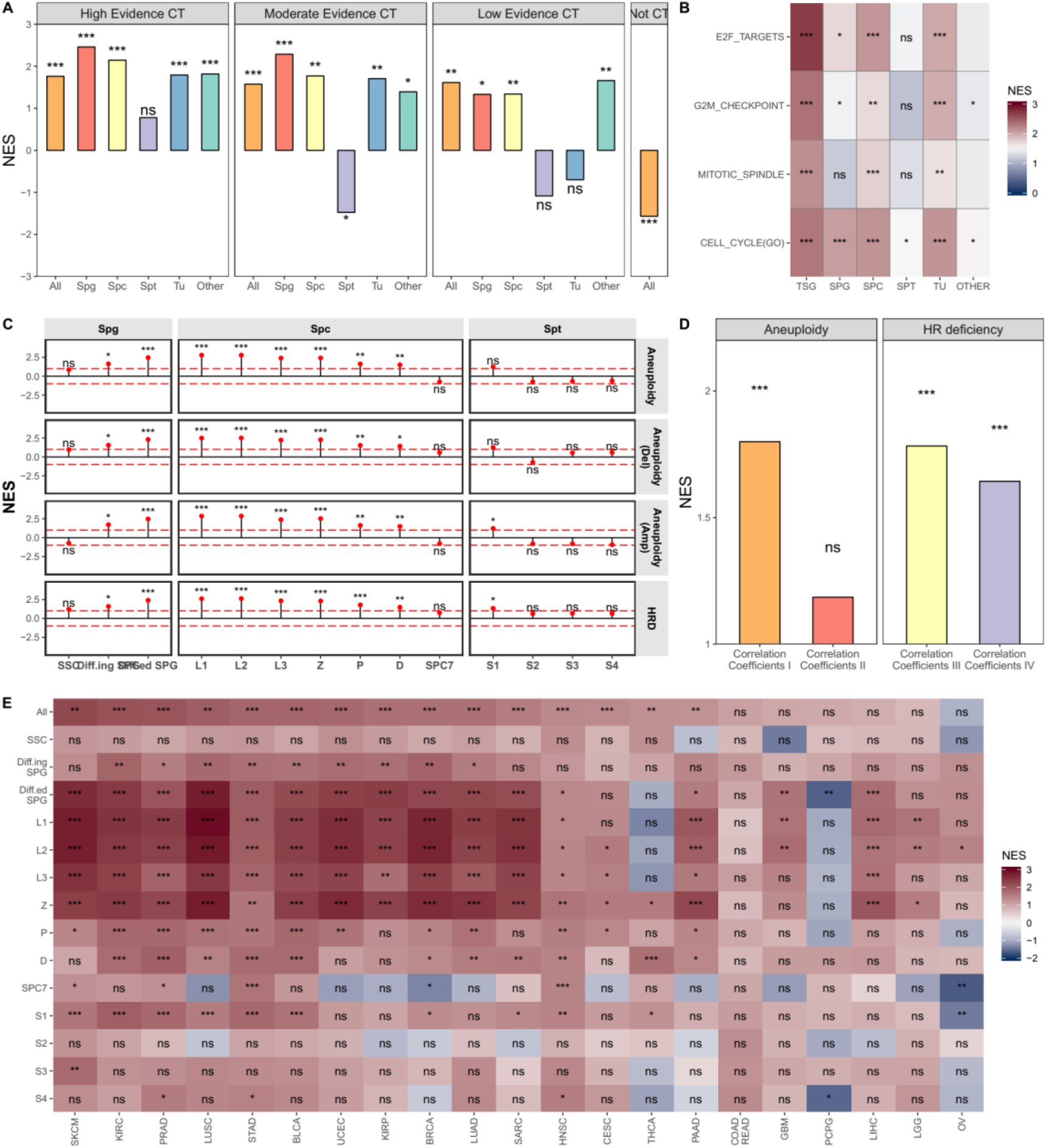
The association between protein-coding CT genes and aneuploidy level in 21 cancer types. ***: P<0.001; **: P<0.01; *: P<0.05 A. Coding CT genes, especially Spg- and Spc-specific expressed CT genes were consistently associated with aneuploidy score; B. The association between CT genes and aneuploidy scores was independent of the proliferation pathways; C. A statistically significant enrichment is observed for the specifically expressed genes in meiosis stages, from differentiated spermatogonia (Diff.ed SPG) to diplotene spermatocytes (D); D. The association between CT genes and HRD were independent of their association with aneuploidy; E. Enrichment analysis was conducted in each tumor types.

A previous TCGA study has revealed that proliferation pathways (such as E2F targets, mitotic spindle, and G2M checkpoint) were positively associated with aneuploidy scores. A number of CT genes were also included in the pathways mentioned above. Thus, we conducted GSEA analysis in each pathway to prove that the association between CT genes and aneuploidy scores was independent of the proliferation pathways. In E2F target gene set, for example, genes with cancer-testis expression pattern showed significantly stronger correlation coefficients with aneuploidy level than the other non-CT E2F target genes (NES = 2.69, *P* = 1.01 × 10^-4^, Figure 2B). Specially, Spc-specific CT genes had a largest normalized enrichment score (NES = 2.10, *P* = 1.10× 10^-4^, Figure 2B). The other proliferation pathways and cell cycle gene set from Gene Ontology showed similar results (Figure 2B).

**Figure 2.**
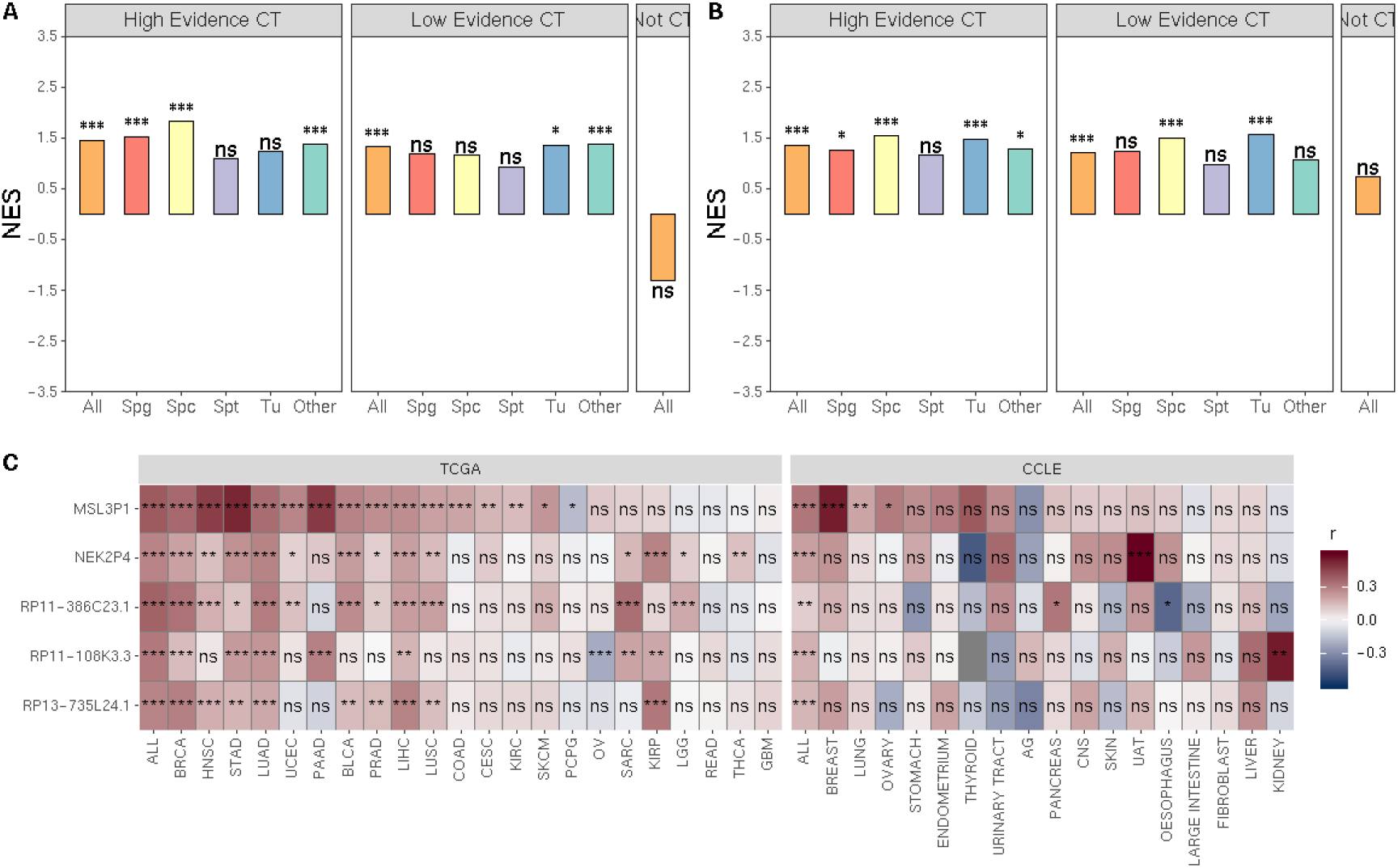
The association between non-coding CT genes and aneuploidy level in 21 cancer types. ***: P<0.001; **: P<0.01; *: P<0.05 A. Non-coding CT genes, especially Spg- and Spc-specific expressed CT genes were consistently associated with aneuploidy score in TCGA data; B. Non-coding CT genes, especially Spg- and Spc-specific expressed CT genes were consistently associated with aneuploidy score in CCLE data; C. Five non-coding CT genes were identified to be associated with aneuploidy in TCGA and CCLE databases.

Next, we further conducted GSEA on specifically expressed genes determined in 14 germ cell subtypes in single-cell RNA-Seq data reported by Wang et al. Interestingly, we observed a statistically significant enrichment for the specifically expressed genes in meiosis stages, from differentiated spermatogonia (Diff.ed SPG) to diplotene spermatocytes (D)(Figure 1C). A similar pattern was seen in the analyses on amplifications and deletions (Figure 1C). During meiosis, programmed double-strand breaks (DSB) formed and are repaired by homologous recombination (HR), which is commonly deficient and lead to the instability of cancer genomes (11). In TCGA patients, the HR deficiency index was highly correlated with aneuploidy score (r=0.63, *P* < 2.20 × 10^-16^). Here, we noticed a consistently positive correlation between meiosis-related CT protein-coding genes and index of HR deficiency. More importantly, if we adjusting the HR deficiency index in the calculation of the partial correlation coefficients between genes and aneuploidy score, the enrichment for CT genes vanished (NES = 1.19, P = 0.17, Figure 1D). In contrary, if we adjusting the aneuploidy score in the calculation of the partial correlation coefficients between genes and HR deficiency index, the enrichment for CT genes are still significant (NES = 1.64, P < 1.00 × 10^-4^, Figure 1D), indicating the correlations between CT genes and aneuploidy were dependent on HR deficiency. Thus, we mainly focused on the correlation between CT genes and HR deficiency in the following analysis. Then, we evaluated the correlation between gene expression and HR deficiency index in 777 CCLE cell lines of solid tumors and successfully validated the enrichment for specific CT genes from meiosis stages (Supplementary Figure 2).

We also performed the same analysis in each TCGA tumor type. We found the significant enrichment for CT genes in the majority of tumor types (15/21, 71.4%) except for nervous system tumors (GBM: Glioblastoma multiforme, LGG: Lower Grade Glioma and PCPG: Pheochromocytoma and paraganglioma), digestive system cancer (COADREAD: Colon and rectal adenocarcinoma, LIHC: Liver Hepatocellular Carcinoma) and Ovarian Cancer (Figure 1E), indicating the different origin of HR deficiency and aneuploidy. Consistent with pancancer analysis, the enriched gene sets mainly involved in the meiosis stages (Figure 1E). Generally, the enrichment scores were higher in early meiosis stages from differentiated spermatogonia to zygotene then in late meiosis stages from pachytene to diplotene (Figure 1D).

### CT non-coding genes were also positively correlated with aneuploidy score in cancers

In our previous work, we noticed that non-coding genes with CT expression pattern also contribute to the development of cancers. Thus, we further explored the association between CT non-coding genes and aneuploidy level in cancers. Because we only obtained expression quantification of 1845 CT genes from single cell RNA-Seq data, we used the CT gene’s classification based on RNA-Seq data from three germ cell subtypes (12). GSEA analysis in TCGA cancer patients revealed the enrichment for high CT non-coding genes (NES = 1.27, *P* = 2.40 × 10^-4^, Figure 2A) and the enrichment was immediately validated in the data from CCLE cell lines (NES = 1.36, *P* = 1.00 × 10^-4^, Figure 2B).

Then, we selected high evidence CT non-coding genes with the top one percent partial correlation coefficients with HR deficiency index for further investigation. A total of 5 CT non-coding genes (MSL3P1, NEK2P4, RP11-386C23.1, RP11-108K3.3, and RP13-735L24.1) were included in the TCGA pancancer analysis and their correlation with HR deficiency were validated in CCLE data (Figure 2C). Among the 5 CT non-coding genes, pseudogene MSL3P1 was the only one detected by testis cell single cell RNA-Seq. It was highly expressed in spermatocytes and the expression was highest in diplotene spermatocytes (Supplementary Figure 3). In the correlation analysis in each tumor type in TCGA and CCLE datasets, we observed that the association between MSL3P1 expression and HR deficiency index were consistently in breast invasive carcinoma (BRCA), lung cancer (lung adenocarcinoma, LUAD; lung squamous carcinoma, LUSC), and ovarian cancer. Because BRCA and OV patients are commonly affected by HR deficiency derived from BRCA1/BRCA2 disruption, we investigated the functional role of MSL3P1 in LUAD and LUSC cell lines. Knockdown of MSL3P1 in A549 cells (a lung adenocarcinoma cell line) and NCI-H1703 cells (an NSCLC squamous carcinomas cell line) could effectively inhibit cell proliferation and clone formation ability (data not show).

### The “off” of RFX2/NFYA and “on” of E2F7/E2F8/MYBL2 altered the regulatory networks of homologous recombination break and repair

Next, we studied why the reactivation of CT genes, which promises the well-organized meiosis in testis, were correlated with aneuploidy in cancers. Here, we first defined 81 CT protein-coding genes as HRD-CT genes, which had the top 1% partial correlation coefficients with HR deficiency index. Then, we constructed a co-expression network by Weighted Correlation Network Analysis (WGCNA) in 259 GTEx testis tissues to explore the relationship between the HRD-related CT protein-coding genes. We identified 3 co-expression modules in normal testis tissues. Interestingly, we found that the majority of CT protein-coding genes (78.7%) and 80 of 81 HRD-related CT genes (98.8%) belonged to Module I (Supplementary Figure 4). In addition, the connectivity level of HRD-CT genes was significantly higher than the other CT genes in Module I (Supplementary Figure 5). Next, we constructed a co-expression network in each TCGA tumor type. Most HRD-CT genes (63.0%-100%) were still classified into a single module, however, a large number of Module I genes (27.3%-98.2%) lost its co-expression pattern with other Module I genes. In addition, the Module I genes was re-classified into multiple modules in tumors (7-34) (Supplementary Figure 6). The results suggested that the co-expression pattern between HRD-CT genes remained in tumors, but the pattern between HRD-CT genes and other Module I CT genes was lost.

The alterations of the co-expression pattern commonly indicated the change of upstream regulators. Thus, we applied Single-Cell Regulatory Network Inference And Clustering (SCENIC) methods (13) to determine which transcription factors (TFs) contribute to the change of co-expression network in testis and cancers. Here, we included the single cell RNA-Seq data from testis and two cancer types (14,15) (Head and Neck squamous cell carcinoma, HNSC; skin cutaneous melanoma, SKCM) to infer the regulatory network. We identified 6 active TFs in testis cells targeting HRD-CT genes, including two groups. In testis cells, Group I (BRCA1, NFYB, E2F1, and TFDP1) were activated in the stages of double-strand breaks during early meiosis and Group II (RFX2 and NFYA) were activated in the stages of double-strand repair during late meiosis (16)(Figure 3A). In HNSC and SKCM single-cell RNA-Seq data, we observed that Group I TFs were still activated in the tumor cells and regulated the HRD-CT genes, but Group II TFs were not activated in cancers(Figure 3B&C). Instead, Group III (E2F7 and E2F8) consistently co-activated with Group I TFs in two types of cancer cells (Figure 3B&C).

**Figure 3.**
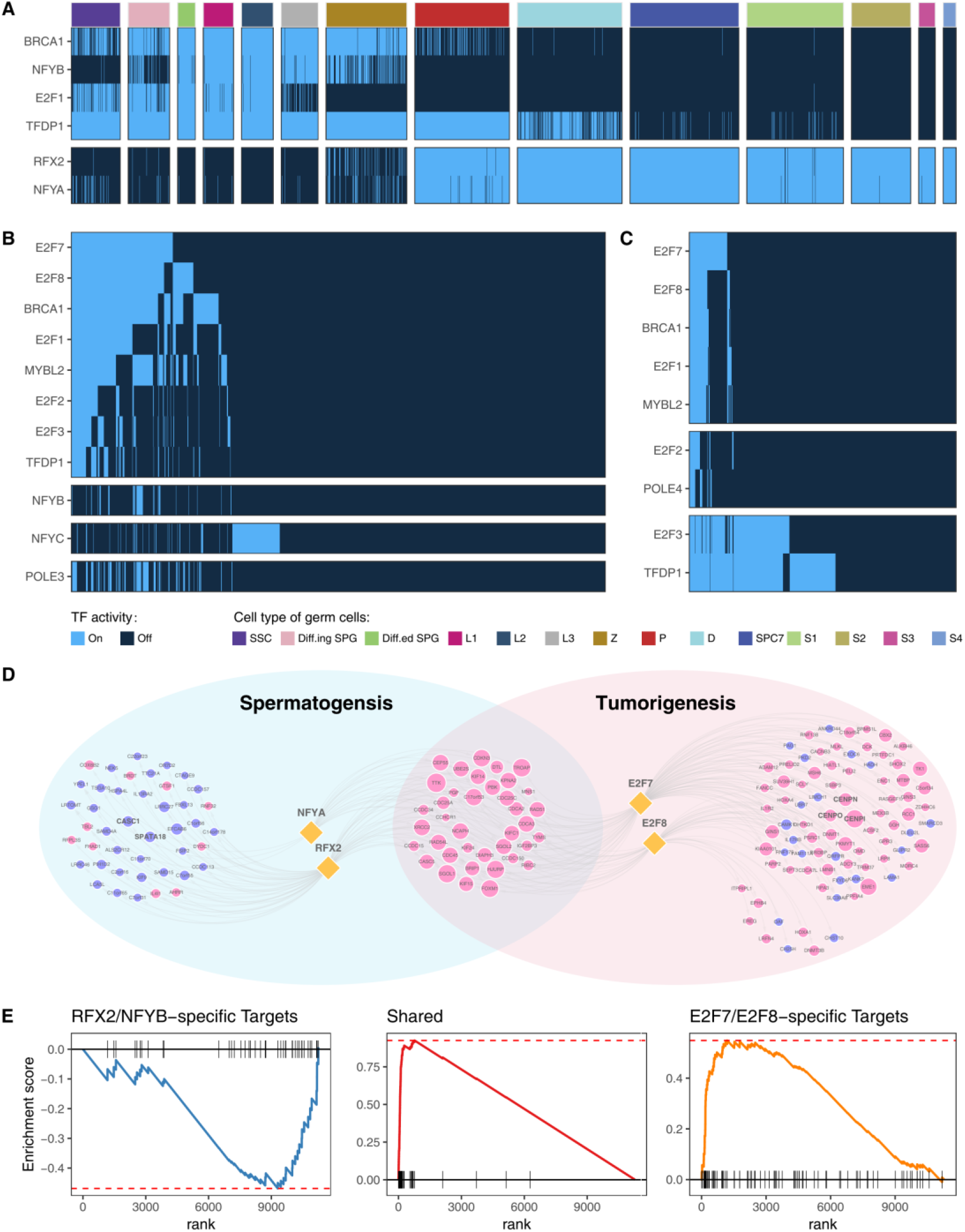
Single-Cell Regulatory Network Inference in testis cells and tumor cells. A. A total of 6 active TFs in testis cells targeting HRD-CT genes. B&C. In HNSC and SKCM single-cell RNA-Seq data, we observed that Group I TFs were still activated in the tumor cells and regulated the HRD-CT genes, but Group II TFs were not activated in cancers. Instead, Group III (E2F7 and E2F8) consistently co-activated with Group I TFs in two types of cancer cells. D&E. We observed a significantly negative enrichment for the CT genes activated only by Group II TFs in testis and a significantly positive enrichment for the CT genes activated only by Group III TFs in cancers.

Moreover, we observed a significantly negative enrichment for the CT genes activated only by Group II TFs in testis (NES=-1.94, P = 4.65 × 10^-4^, Figure 3D&E) and a significantly positive enrichment for the CT genes activated only by Group III TFs in cancers (NES=2.18, P = 1.15 × 10^-3^, Figure 3D&E). Given the above observations, we conclude that the sequential activation of Group I and Group II TFs during the meiosis may promise the balanced double-strand break and repair respectively, while the gain of Group III TFs strengthened double-strand break related with Group I TFs and the loss of Group II TFs weakened the repair process, which contributes to the homologous recombination deficiency and aneuploidy in cancers.

### The correlation between CT genes and HRD diminished their correlation with infiltrating cytotoxic immune cells

Recent studies found that highly aneuploid tumors presented reduced expression of markers of cytotoxic infiltrating immune cells, especially CD8+ T cells. However, CT proteins were famous for their immunogenicity, which can induce the immune response. Thus, we further investigate whether the correlation between CT genes and immune infiltrating cells in cancers could be influenced by their association with HRD. In the analysis of the correlation between CT genes and infiltrating CD8+ T cells, we observe a significant enrichment for Spc-specific CT genes (NES=1.51, P = 2.14 × 10^-3^). Interestingly, the enrichment score for Spg- and Spc-specific CT genes increased greatly after adjusting the HRD index in the calculation of partial correlation coefficients (NES_Spg_=1.94, P = 7.29 × 10^-4^; NES_Spc_=1.99, P = 7.29 × 10^-4^, Figure 4A). No enrichment was found for the Spt-specific and universally expressed CT genes. A similar increase was also seen in the analysis of correlation enrichment between Spg-specific CT genes and NK cells (NES_model1_=1.50, P = 4.70 × 10^-3^; NES_model2_=2.50, P = 9.27 × 10^-3^, Figure 4A). But for the other infiltrating lymphocytes, we did not found such an increase.

**Figure 4.**
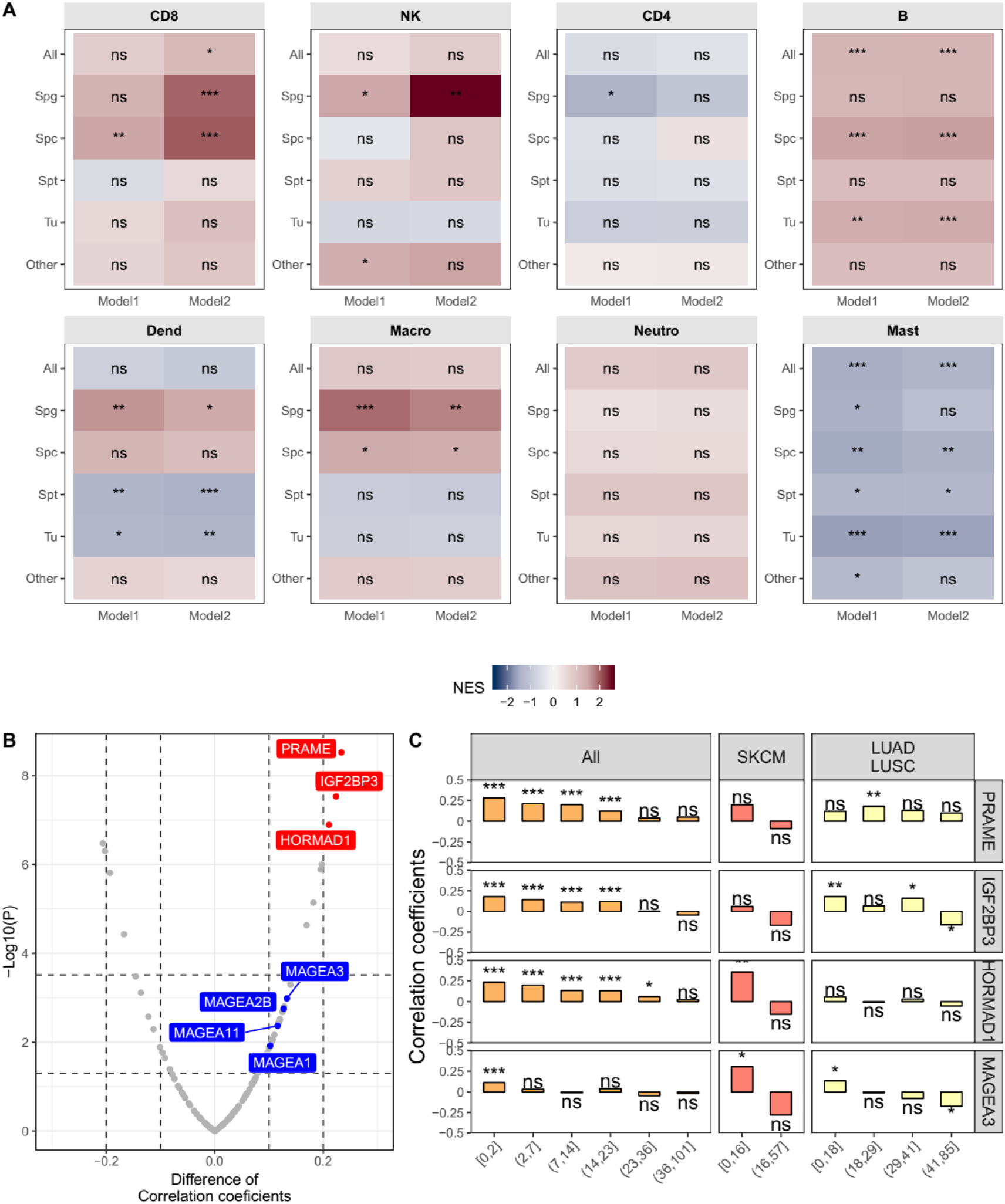
The correlation between CT genes and immune infiltrating cells in cancers was influenced by their association with HRD. A. For CD8+ T cells, the enrichment score for Spg- and Spc-specific CT genes increased greatly after adjusting the HRD index in the calculation of partial correlation coefficients; B. Three CT antigens (PRAME, IGF2BP3, and HORMAD1) was identified to correlate with CD8+ T cells only in HRD low patients; C. In addition to the three genes, the correlation between MAGEA3 and CD8+ infiltrating cells also significantly decreased in the patients with high HRD level

To further investigate which CT antigens’ immunogenicity could be influenced by the HRD level, we compared the partial correlation coefficients between previously defined CT antigens and CD8+ infiltrating cells across the patients with lowest HRD score (0-2) and those with highest HRD score (36-101). Three CT antigens (PRAME, IGF2BP3, and HORMAD1) was identified after Bonferroni correction (0.05/127) and showed higher correlation coefficients in patients with lowest HRD level than in patients with highest HRD level (Figure 4B). Further analyses revealed that the correlation coefficients between the expression of three CT antigens and the level of CD8+ infiltrating cells decreased with the HRD score increasing (Figure 4C). In patients with SKCM and Pan-lung cancer (LUAD and LUSC), similar trends were also observed. MAGEA3 is a member of melanoma antigen family A genes and its ability to elicit spontaneous humoral and cellular immune responses have been shown in cancer patients. However, recent clinical trials did not support its benefit as a target of immunotherapy. Here, we noticed that the correlation between MAGEA3 and CD8+ infiltrating cells also significantly decreased in the patients with high HRD level (P=0.001, Figure 4B&C) though the difference did not pass the rigor of Bonferroni correction, suggesting that the HRD level may be responsible for the failure of immunotherapy targeting MAGEA3.

## Discussion

The processes of germ cell development and tumor development share interestingly similarities. CT genes, which are specifically expressed in testis and cancer tissues, were regarded as the molecular basis of the two processes. However, spermatogenesis is a tightly regulated developmental process and include multiple phases(10). In each phase of human spermatogenesis, various developmental germ cell subtypes can be identified. Thus, systematic investigation of the CT genes expressed in specific germ cell subtypes will help the understanding of the similarity between germ cells and cancers. However, the dynamic transcriptome of the developmental process of sperm is difficult. Recently, the application of single-cell RNA-Seq on testicular cells greatly extended our knowledge on spermatogenesis and made it available to identify the spermatogenesis stage(s) similar to cancer development. In the present study, we integrated multi-omics data from bulk samples and single cell RNA-Seq data to prove that CT genes involved in the meiosis stages were consistently correlated with aneuploidy in cancers. Although the aneuploidy-related CT genes were activated in the testis cells as well as in the cancer cells, our data showed their activation transcriptional factors were different. In addition, we found that the correlation between CT genes and aneuploidy can influence the immunogenicity of CT antigens. Detailed information concerning the CT genes and their association with aneuploidy and tumor lymphocytes infiltration in this article is freely accessible through our search engine (CTatlas2: http://45.62.103.54:41062/ctatlas2/).

However, meiosis is a unique and specialized cellular division process to create haploid gametes. Homologous recombination is a hallmark of meiosis. In order to initiate homologous recombination and ensure proper homologous chromosome segregation, cells deliberately form numerous DNA double-strand breaks, though they are highly hazardous for genome integrity(17) Thus, meiotic DSB formation, processing, and repair are accurately regulated to promise their occurrence at the right time and place. Although the activation of CT genes can trigger the alteration of ploidy in the process of spermatogenesis as well as tumorigenesis, the process of aneuploidy generation accompanied chaotic breaks of DNA, which is quite distinct from the well-organized haploid sperm cells generation. Our results revealed a group of “accelerator” transcriptional factors, such as E2F7 and E2F8, can activate the genes associated with HRD as well as a series of genes involved in the cell cycle and proliferation. The overexpression E2F7 have been reported to result in an increase of aneuploidy in breast cancer cells(18). On the contrary, another group of “stabilizer” transcriptional factors, such as RFX2 and NFYA, can activate a number of CT genes to promise the timely repair of double-strand breaks in the late meiosis stages, but these CT genes were not co-activated with HRD related CT genes in cancer cells. Thus, the gain of “accelerator” and the loss of “stabilizer” may contribute to the aberrant aneuploidy in cancers. These results also suggested that CT genes can serve as aneuploidy suppressors, though they were considered as oncogenes in cancers. CASC5 and SPATA18 represented these suppressors, as they were negatively correlated with the aneuploidy and HRD level and the knock out of the genes can result in the development of cancers (19).

Another important point should be mentioned is that the association between CT genes and aneuploidy may influence the immunogenicity of CT genes. CT genes were first observed and considered to be candidate targets for immunotherapy due to their immunogenicity. However, 19 of the 24 phase II/III clinical trials that tested 17 distinct therapeutic anticancer vaccines failed to achieve their primary objectives(20–23); this included a trial for the vaccine targeting MAGEA3 in non-small cell lung cancer (NSCLC) and melanoma. A recent study reported that the aneuploidy level can predict the reduced level of cytotoxic infiltrating immune cells (8). Our results further suggested that the association between CT genes and cytotoxic infiltrating immune cells dependent on the aneuploidy level, including famous MAGEA3. The results suggested that the immunotherapy targeting on CT antigens should carefully evaluate their association with aneuploidy.

## Methods

### Public databases

We included genomic and transcriptomic data from bulk samples from 8,879 TCGA patients with solid tumors, and 777 cancer cell lines from CCLE databases. We also included single cell RNA-Seq data by Smart-Seq2 Technology from testis cells and two cancer types. The databases and related information were provided in Supplementary Table 1. The abbreviation of cancer and sample sizes from the TCGA project were provided in Supplementary Table 2.

### RNA-Seq quantification

For RNA-Seq data from three germ cell subtypes from Jan et al.’s study, we applied standard expression quantification pipelines. Briefly, RNA reads were generated, aligned to the GENCODE v19 genome assembly with STAR v2.4.148, and quantified with RSEM (RSEM v1.2.12). TPM was used for the quantification of gene expression and following analysis.

### Homologous recombination deficiency index

We calculated the homologous recombination deficiency index by an R package scarHRD, which determines the levels of homologous recombination deficiency (telomeric allelic imbalance, loss of heterozygosity, number of large-scale transitions) based on an array or sequencing data.

### CT genes and the further classification of CT genes

We obtained CT genes information from our previous work(2). CT genes were classified into the following five categories based on specificity measure (SPM) values:

1. Spermatogonia (Spg) specific CT genes (SPM_Spg_>0.9 and expression in Spg > 0.1);
2. Spermatocyte (Spc) specific CT genes (SPM_Spc_>0.9 and expression in Spc > 0.1);
3. Spermatids (Spt) specific CT genes (SPM_Spt_>0.9 and expression in Spt > 0.1);
4. Testis-universal (Tu) expressed CT genes (SPM_Spg_<0.9, SPM_Spc_<0.9 and SPM_Spt_<0.9 and expression in Spg, Spc and Spt > 0.1)
5. Other CT genes.

Further classification of 14 subtypes based on the differential expressed genes provided by the study of Wang et al.

### Gene set enrichment analysis (GSEA)

A gene set enrichment analysis (GSEA) was performed using partial Spearman’s rank correlation coefficients between the all genes and aneuploidy level, HRD and infiltrating lymphocytes using the “fgsea” command from the Bioconductor package fgsea based on the CT genes classification mentioned above.

### Construction of the regulatory network

The SCENIC analysis was run as described on the cells that passed the filtering, using the 20-thousand motifs database for RcisTarget and GRNboost (SCENIC version 0.1.5, which corresponds to RcisTarget 0.99.0 and AUCell 0.99.5; with RcisTarget.hg19.motifDatabases.20k). The input matrix was the TPM for each dataset. Only shared genes across the databases were used for network construction. Only germ cells and tumor cells were used for analyses.

### Statistic methods

All statistical tests were performed using a Wilcoxon rank-sum test for continuous data, Spearman’s rank correlation or partial correlation for the estimation of correlation. Fisher’s exact test was used to assess differences for the count data. Multiple testing corrections were performed where necessary using the Benjamini-Hochberg method. All reported *P* values are two-sided. Figures were generated with the R packages ggplot2(24) and RColorBrewer(25).

---

**Supplementary Figure 1.**
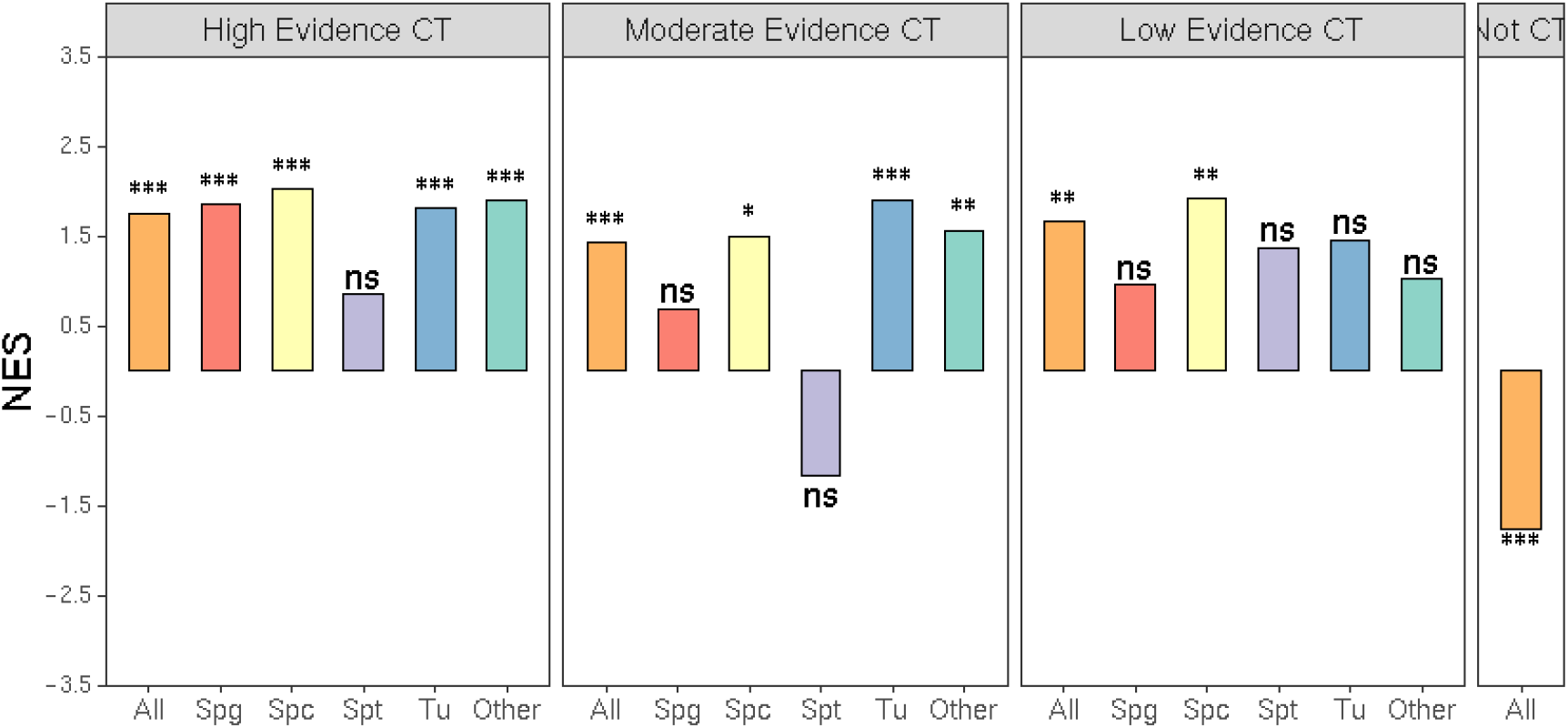
An independent RNA-Seq data which include the transcriptomic profile of cells from three well-defined germ cell subtypes in testis validated the classification and the enrichment.

**Supplementary Figure 2.**
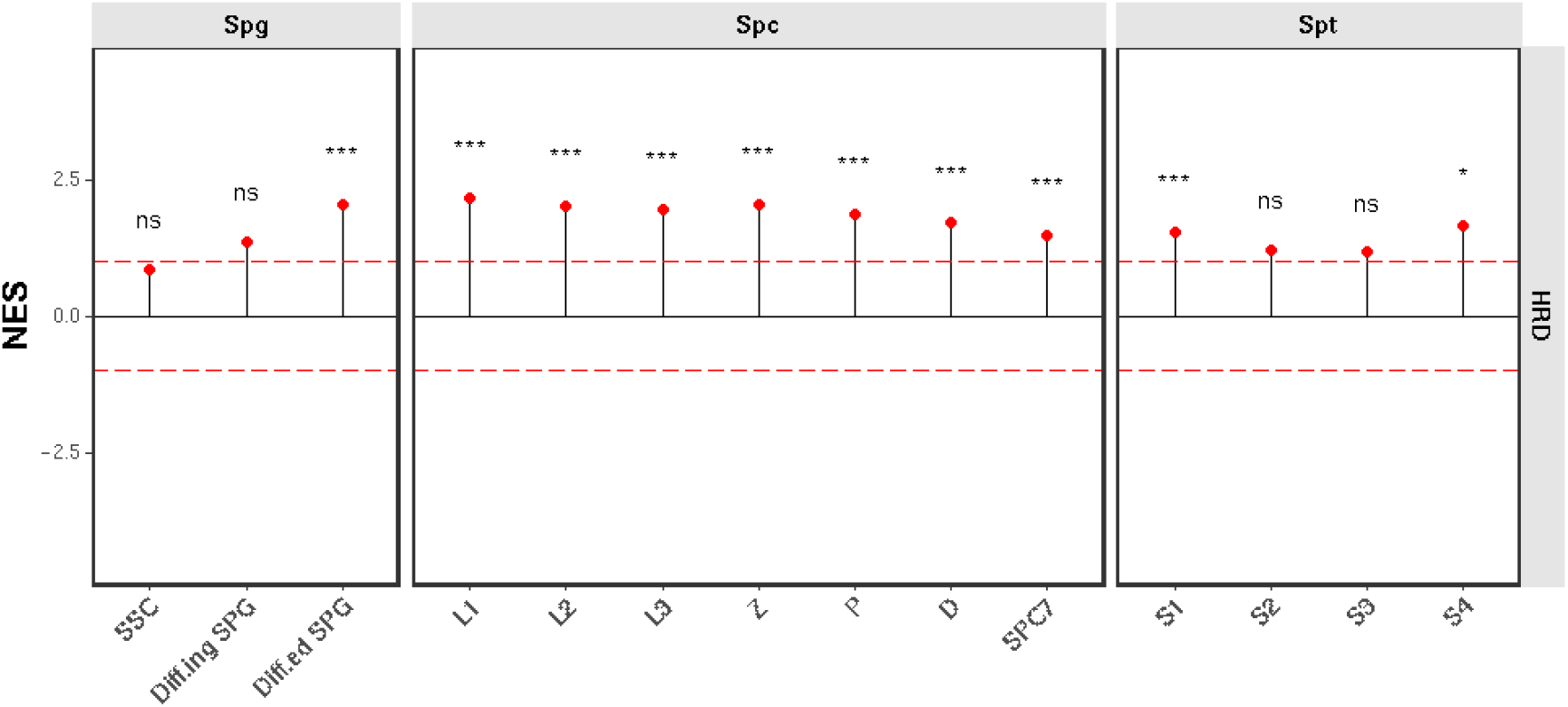
The correlation between gene expression and HR deficiency index in 777 CCLE cell lines of solid tumors and the enrichment analysis.

**Supplementary Figure 3.**
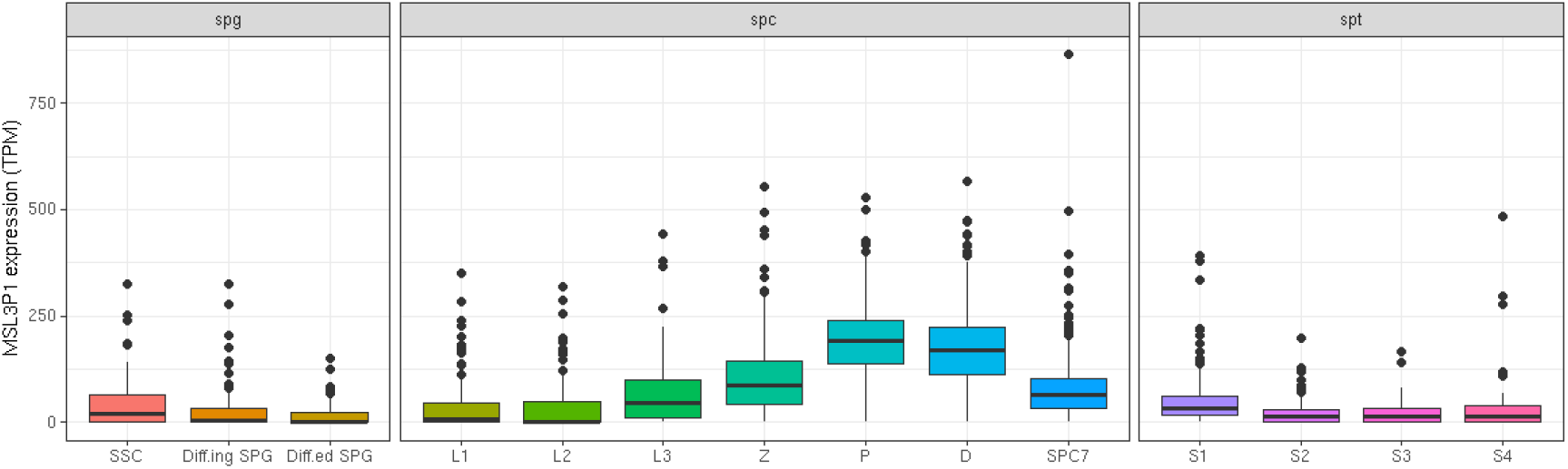
The expression pattern of MSL3P1 in the different stages of testis cells.

**Supplementary Figure 4.**
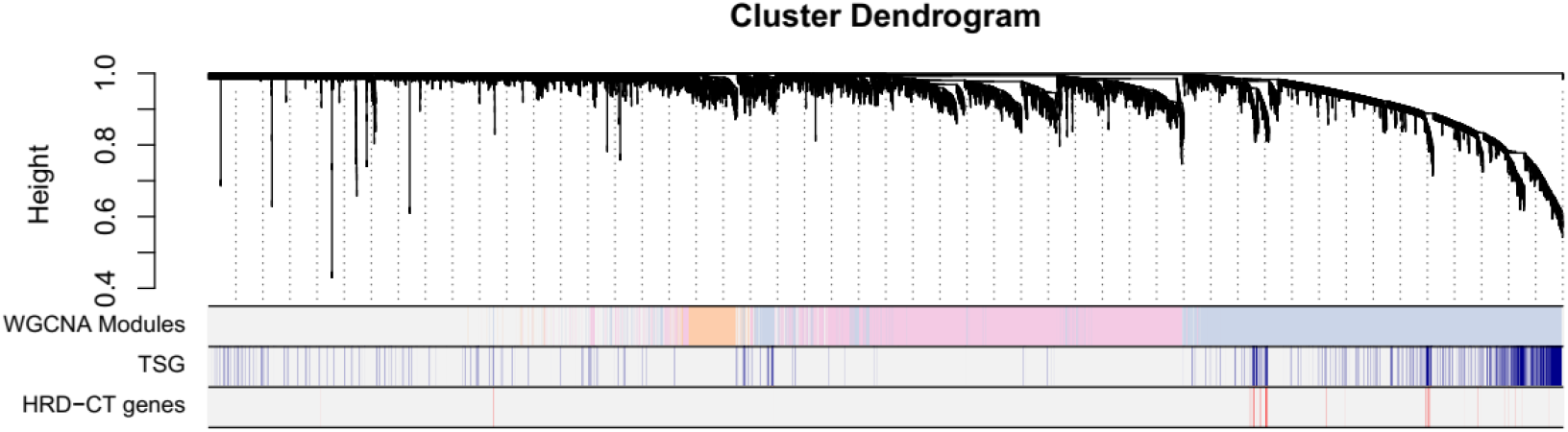
WGNCA constructed the co-expression network in testis.

**Supplementary Figure 5.**
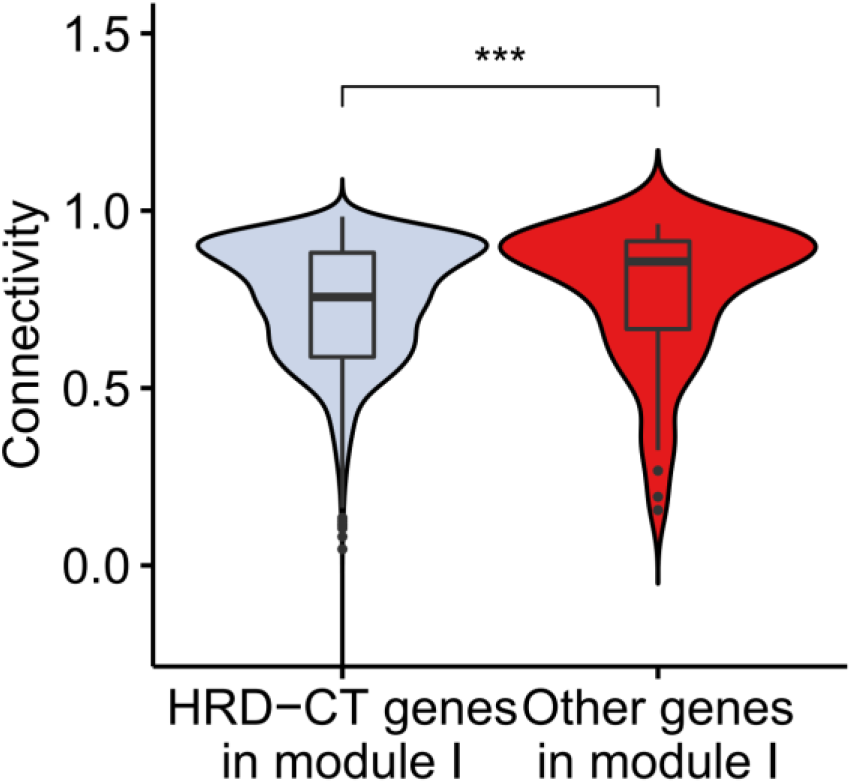
The connectivity level of HRD-CT genes was significantly higher than the other CT genes in Module I.

**Supplementary Figure 6.**
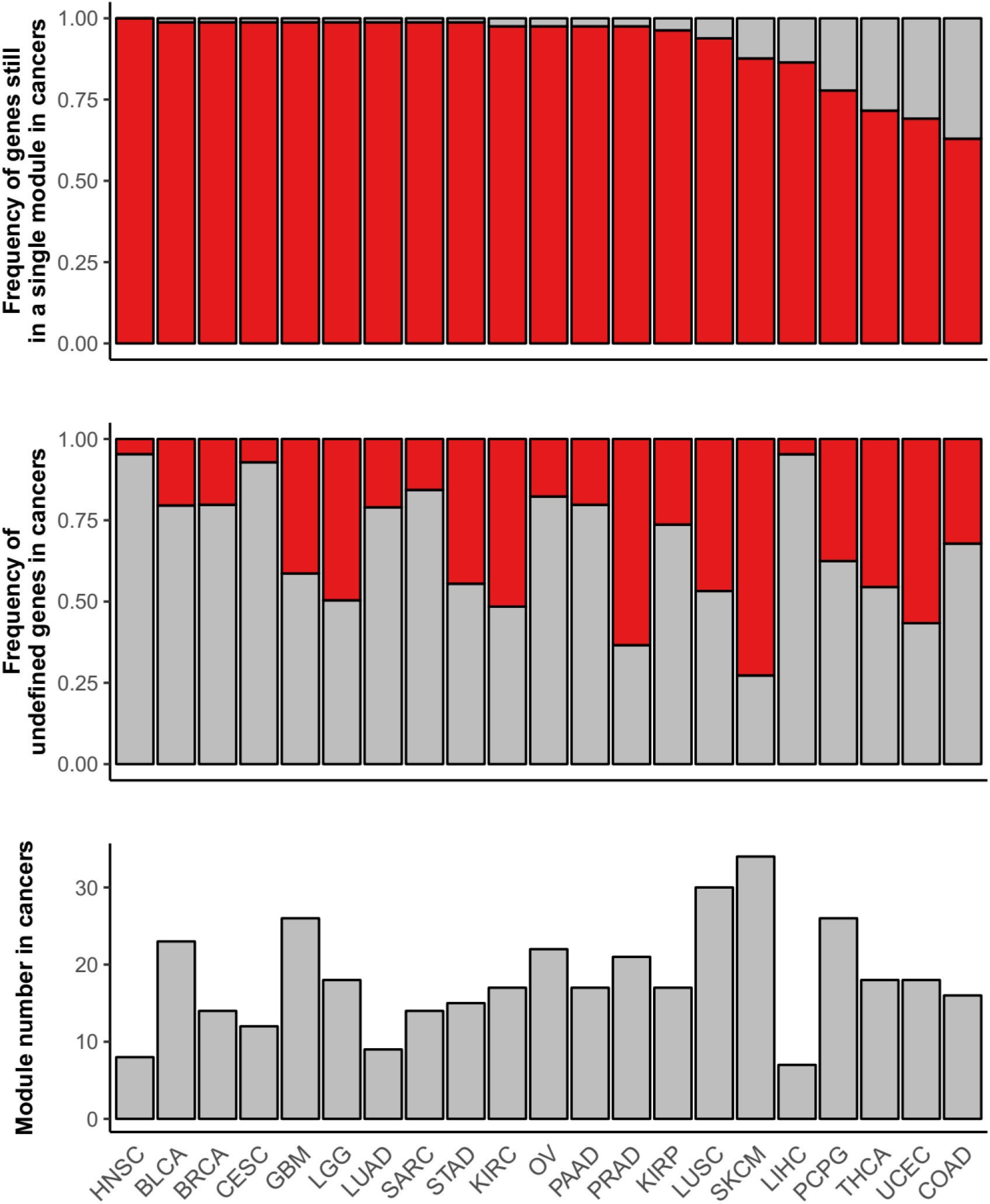
Most HRD-CT genes (63.0%-100%) were still classified into a single module, however, a large number of Module I genes (27.3%-98.2%) lost its co-expression pattern with other Module I genes.

**Supplementary Table 1.**
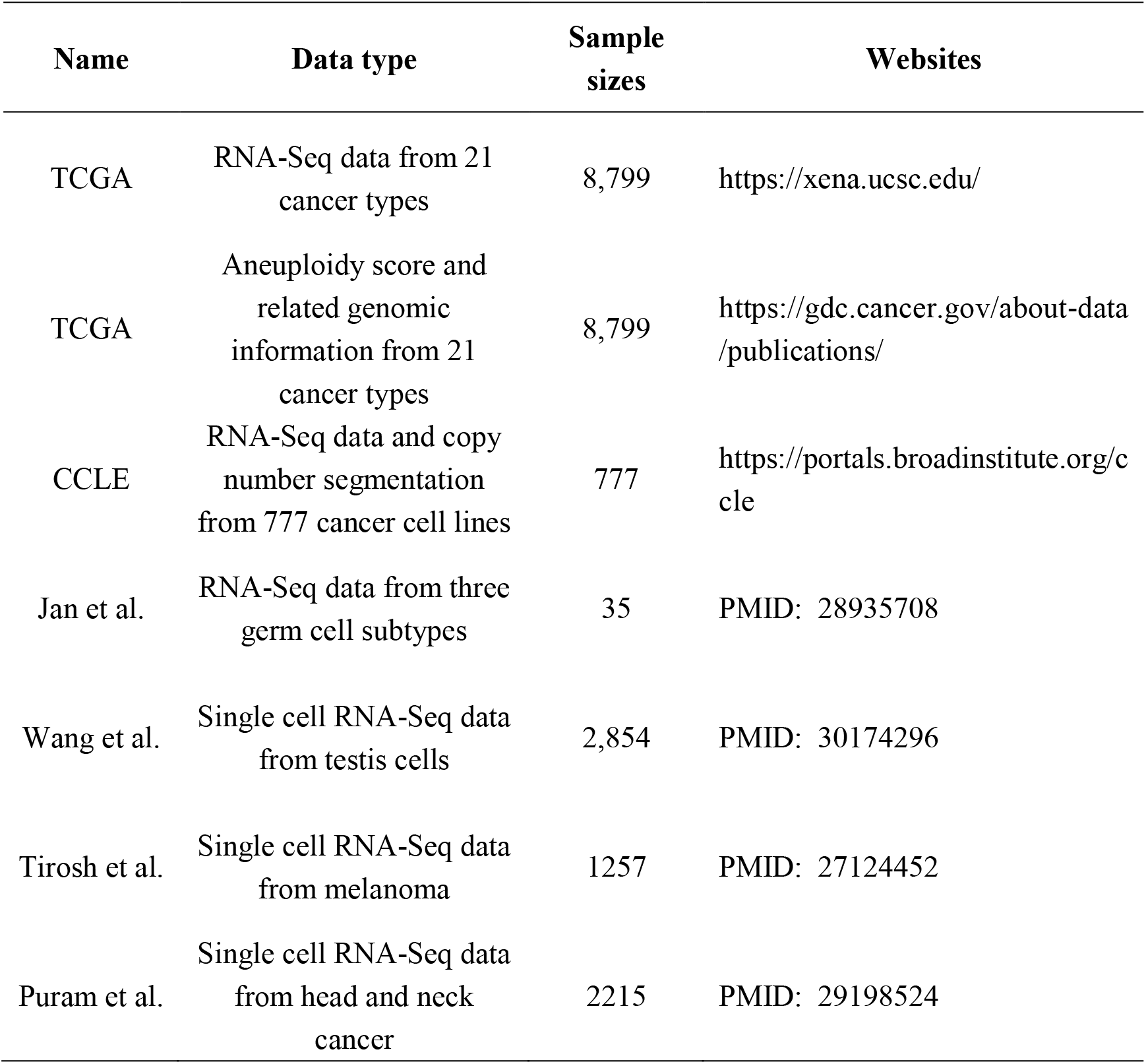
The databases and related information

**Supplementary Table 2.** The abbreviation of cancer and sample sizes from the TCGA project

| **Tumor** | **Full name** | **Sample size** |
| --- | --- | --- |
| BRCA | Breast invasive carcinoma | 1,091 |
| COAD | Colon adenocarcinoma | 622 |
| UCEC | Uterine Corpus Endometrial Carcinoma | 543 |
| KIRC | Kidney renal clear cell carcinoma | 530 |
| LUAD | Lung adenocarcinoma | 513 |
| LGG | Brain Lower Grade Glioma | 511 |
| THCA | Thyroid carcinoma | 502 |
| LUSC | Lung squamous cell carcinoma | 501 |
| HNSC | Head and Neck squamous cell carcinoma | 500 |
| PRAD | Prostate adenocarcinoma | 495 |
| BLCA | Bladder Urothelial Carcinoma | 408 |
| STAD | Stomach adenocarcinoma | 375 |
| OV | Ovarian serous cystadenocarcinoma | 374 |
| LIHC | Liver hepatocellular carcinoma | 371 |
| CESC | Cervical squamous cell carcinoma and endocervical adenocarcinoma | 304 |
| KIRP | Kidney renal papillary cell carcinoma | 288 |
| SARC | Sarcoma | 259 |
| PCPG | Pheochromocytoma and Paraganglioma | 178 |
| PAAD | Pancreatic adenocarcinoma | 177 |
| GBM | Glioblastoma multiforme | 154 |
| SKCM | Skin Cutaneous Melanoma | 103 |

